# A rapid, accurate approach to inferring pedigrees in endogamous populations

**DOI:** 10.1101/2020.02.25.965376

**Authors:** Cole M. Williams, Brooke A. Scelza, Sarah D. Slack, Rasika A. Mathias, Harold Watson, Kathleen C. Barnes, Ethan Lange, Randi K. Johnson, Christopher R. Gignoux, Sohini Ramachandran, Brenna M. Henn

## Abstract

Accurate reconstruction of pedigrees from genetic data remains a challenging problem. Pedigree inference algorithms are often trained only on European-descent families in urban locations. Many relationship categories can be difficult to distinguish (e.g. half-sibships versus avuncular) without external information. Furthermore, existing methods perform poorly in endogamous populations for which there may be reticulations within the pedigrees and elevated haplotype sharing. We present a simple, rapid algorithm which initially uses only high-confidence first-degree relationships to seed a machine learning step based on summary statistics of identity-by-descent (IBD) sharing. One of these statistics, our “haplotype score”, is novel and can be used to: (1) distinguish half-sibling pairs from avuncular or grandparent-grandchildren pairs; and (2) assign individuals to ancestor versus descendant generation. We test our approach in a sample of 700 individuals from northern Namibia, sampled from an endogamous population called the Himba. Due to a culture of concurrent relationships in the Himba, there is a high proportion of half-sibships. We accurately identify first through fourth-degree relationships and distinguish between various second-degree relationships: half-sibships, avuncular pairs, and grandparent-grandchildren. We further validate our approach in a second diverse African-descent dataset, the Barbados Asthma Genetics Study (BAGS). Accurate reconstruction of pedigrees holds promise for tracing allele frequency trajectories, improved phasing and other population genomic questions.

## INTRODUCTION

Geneticists have long relied on pedigrees to map disease-causing alleles, uncover modes of trait inheritance, and to construct recombination maps (Hall et al., 1990; Ott, 1974; Griffiths and Marjoram, 1996; Thompson, 1981; Kong et al., 2002). Traditionally, pedigrees for biomedical studies are constructed using self-reported information from study participants, as in the San Antonio Mexican American Family Studies pedigree (Hunt et al., 2005). Statistical genetics has shifted focus from pedigree studies to genome-wide association (GWA) studies over the past two decades, and GWA studies either prune close-degree relatives or control for relatedness using a kinship matrix (Manichaikul et al., 2010; Eu-ahsunthornwattana et al., 2014; Zaitlen et al., 2013). There is recent renewed interest in genetic relatives and methods to detect them in genomic datasets. Downstream studies enabled by detecting genetic relatives range from estimation of heritability (Zaitlen et al., 2013); to parent-of-origin effects; (Hofmeister et al., 2023; Zeng et al., 2019); to indirect genetic effects (Young et al., 2019, 2018); and inference of kinship and mating patterns (Swinford et al., 2022; Scelza et al., 2020). There is also interest in genetic relatives in both the direct-to-consumer genetic testing space (Jewett et al., 2021) and the emerging field of forensic genealogy (Phillips, 2018; Moon, 2014). As genomic datasets grow in size, the number of close relatives grows quadratically, and so too the need to accurately identify and characterize these close relatives (Ramstetter et al., 2017; Henn et al., 2012).

Motivated by this range of challenges, a number of kinship inference methods have been developed that use various summaries of identity-by-descent (IBD) sharing between relatives to either build pedigrees or identify relative pairs (Staples et al., 2014; Ko and Nielsen, 2017; Manichaikul et al., 2010; Jewett et al., 2021; Qiao et al., 2021; Huff et al., 2011; Li et al., 2014). Much focus has been on what Gimelfarb (1981) calls Type I and Type II relationships; Type I relatives are connected in the pedigree through a single parent (e.g., half-siblings, first cousins) and Type II relatives are connected through two parents of one individual and one parent of the other individual (e.g., an avuncular pair). Type III relatives are both connected through both parents (a classical example are double-cousins, who are first cousins through their mothers and their fathers). The expected amount of IBD shared by two relatives can be easily computed given the length of the path (or *paths*, for Type II and III relationships) that connect the pair through the pedigree. Under assumptions of random mating and infinitely large populations, inferring the degree of relatedness of two individuals using IBD is straightforward, limited only by exponential decay of IBD sharing as pedigree paths get longer (Hill and White, 2013). However, existing methods fail to account for (1) non-traditional (by Western standards) family structures and mating patterns, and (2) bottlenecks, isolation, and/or endogamy, which can skew or shift distributions of IBD sharing (Palamara et al., 2012; Browning and Browning, 2015; Huff et al., 2011). Additionally, existing methods do not model Type III relationships.

Standard pedigree inference methods often ignore endogamy and consanguinity in their discussions of IBD segments and relationship inference. We note that endogamy and consanguinity are distinct but not mutually exclusive. Endogamy is the practice of marrying within a cultural, ethnic, or religious group but does not necessarily occur between close relatives (Bittles and Black, 2010). Endogamy results in increased background IBD sharing, such that the number, length, or position of IBD segments may not reflect the expectation from an ‘outbred’ population (Hill and White, 2013). However, the performance of IBD-based relationship inference software on endogamous or founder populations has rarely been studied. This affects numerous populations worldwide, and is critical to studies of large structured populations, as the norm in multiple geographic regions such as the Near East, South Asia, and Latin America (Nakatsuka et al., 2017; Mooney et al., 2018).

Here, we introduce PONDEROSA, **P**arent **O**ffspri**N**g pe**D**igree inf**E**rence **RO**bu**S**t to endog**A**my, a software package for identifying and characterizing genetic relatives. PONDEROSA traces parent-offspring lineages to find high-confidence relationship pairs and uses these pairs to learn the population distribution of various IBD summary statistics. PONDEROSA is able to classify degrees of relatedness (e.g., 2nd, 3rd, 4th degree relatives) with higher accuracy in datasets with elevated background IBD than existing algorithms. PONDEROSA also employs a novel algorithm for inferring the pedigree relationship (grandparent-grandchild, avuncular or half-sibling) of second degree relatives, including distinguishing between maternal vs. paternal affiliations. The algorithm trains two linear discriminant analysis (LDA) classifiers on canonical properties of IBD sharing (the number of IBD segments and the total lengths of segments) and a novel statistic called the “haplotype score.” The haplotype score can also be used to identify the genetically older individual of a pair (e.g., the parent of a parent-offspring pair or the uncle/aunt of an avuncular pair) without need for age data. We apply PONDEROSA to characterize relatives in two datasets: the endogamous Himba of northern Namibia and another African-descent dataset from Barbados (Barbados Asthma Genetics Study or BAGS). We further illustrate the ability of PONDEROSA and our simulation pipeline to classify complex reticulated pedigree relationships by simulating Gimelfarb’s Type III relationships. We classify these relationships using real genotype data from the Himba and BAGS datasets as well as using synthetic genotype data from 10,000 Quebecois individuals from Anderson-Trocmé et al. (2023). The latter dataset represents a European-descent founder population, the French Quebec, that may have elevated identity-by-descent sharing and dense relatedness. Our simulation results show that Type III relationships may be ubiquitous globally, as we show evidence of these relative pairs in each of these three global populations.

## METHODS

### Generating training data

PONDEROSA generates training data from known pairs in the dataset as outlined in Figure 1. The first phase of PONDEROSA is dedicated to (1) finding or simulating these relative pairs and (2) computing various summary statistics on IBD segment data. An input of PONDEROSA is a PLINK-formatted .fam file, which contains information about the parent-offspring relationships contained in the dataset. We use this information to create a pedigree graph, a directed graph in which edges point from parent to offspring. PONDEROSA first attempts to resolve ambiguous sibships, pairs who share one parent but who are both missing the other parent and could be either half-siblings or full siblings. In an outbred population, we could easily make this decision by looking at the proportion of the genome that is shared IBD1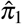 sites are IBD1 if the pair are IBD on one of the two haplotypes) and the proportion of the genome that is IBD2 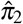 sites are IBD2 if the pair are IBD on both haplotypes). For half-siblings, 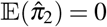 and for full-siblings IE 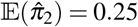. In the Himba, we observed half-siblings with high IBD2 proportions; in order to distinguish half-siblings from full-siblings, we fit a 2-component Gaussian Mixture model with initial means equaling the expectations of IBD1 and IBD2 proportions in an outbred population; note that we do not use the LDA classifier here as we do not yet have the labels. Ambiguous siblings inferred to be full-siblings are given a placeholder parent in the pedigree graph.

**Figure 1.**
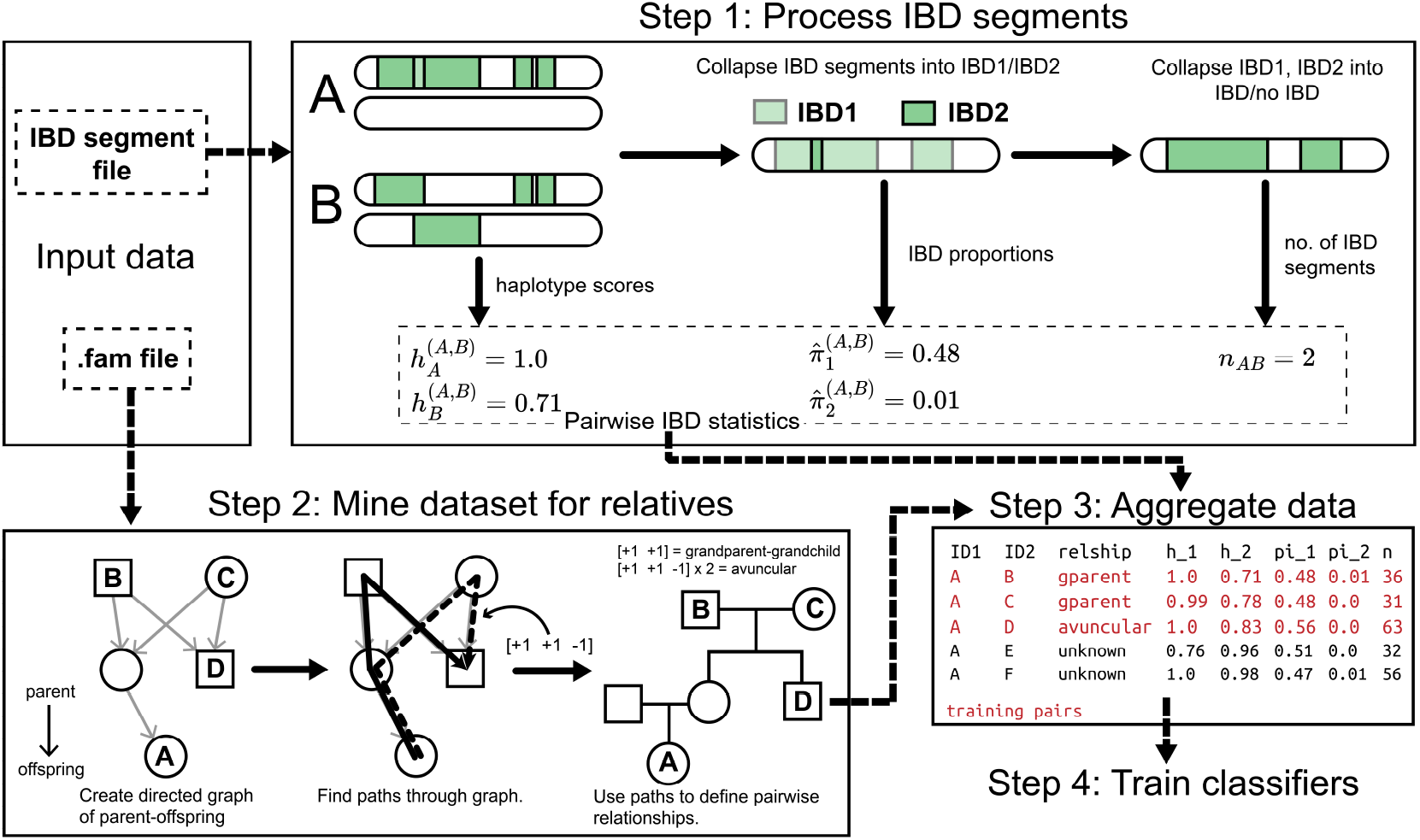
An overview of PONDEROSA. The minimum input data are (1) an IBD segment file that contains pairwise IBD segments, including the start and end genetic coordinate and the haplotype indices corresponding to the two individuals sharing the segment and (2) a PLINK-formatted FAM file containing parent-offspring information. In Step 1, the IBD segments are processed for each pair of relatives, giving five different IBD statistics for each pair. We provide an example of the IBD sharing statistics on a single chromosome for the relative pair (A, B) who are a grandparent-grandchild pair. In Step 2, PONDEROSA builds a parent-offspring graph, a directed graph in which edges direct parents to offspring. PONDEROSA finds paths through the graph, as outlined in Methods. A path is a vector of [+1, -1], describing the movement from node to node (+1 describes moving from offspring to parent and -1 describes moving from parent to offspring). The pair (A,B), for instance, are connected by the path [+1, +1], which is the path for a grandparent-grandchild pair. In Step 3, we merge the IBD sharing statistics with the relationships found in Step 2. Pairs with a known relationship (shown in red) are used to train three LDA classifiers (Step 4).

For a given focal individual, we use a recursive algorithm to find all possible relatives. A path between relatives *i* and *j* is represented as a vector describing the direction of the path: +1 describes a movement “up” the pedigree (from child to parent) and -1 describes a movement “down” the pedigree (from parent to child). For example, a pair of first cousins are described as having two of the following paths: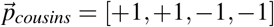. In practice, we also keep track of the parent that starts the path (+1 refers to a paternal path and +2 refers to a maternal path): if two 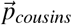 paths go through the same parent, the pair are first-cousins, but if they go through different parents, the pair are double half-cousins. This is an important distinction to make: even though first-cousins and double half-cousins have the same expected kinship coefficient, double half-cousins have an 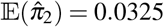 whereas first cousins have an 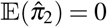. There are two path constraints in the algorithm: (1) paths must start by ascending and (2) once a path descends, it cannot ascend. PONDEROSA uses a set of pre-defined relationships that are described by their paths through the pedigree. The benefit to finding relatives in this way is that it is not rule-based and allows the user to easily supply additional relationships that are not pre-defined in PONDEROSA, e.g., PONDEROSA does not automatically look for great-avuncular relationships but the user can easily add the relationship.

Finding enough high-confidence pedigree relationships in the dataset may be limited in small datasets with few parent-offspring pairs. To overcome this limitation, we developed a pipeline for simulating relative pairs using Ped-sim (Caballero et al., 2019). To simulate a paternal half-sibling pair, for example, Ped-sim would require three founders: a father and the two mothers. To simulate 100 paternal half-sibling pairs, the input VCF would need 300 individuals. For each simulation, Ped-sim randomly chooses three individuals as founders; this random choice means that two close relatives could be chosen as founders. To overcome this, the pipeline will reject a simulation if, out of all founder pairs, there is at least one pair that is more related than a user-specified kinship coefficient. This can be set by looking at mating pairs in the dataset or estimated through the distribution of autozygosity in the dataset. In the initialization step, we sample 10,000 sets of founders to compute the probability of rejecting a simulation and estimate the number of simulations we will need to run to achieve the desired number of training pairs. For example, if the probability of rejecting a simulation is 40% and we want 100 pairs, we would need to simulate 182 pairs in order to have a 90% chance of accepting at least 100 simulations. Depending on how many samples are in the input VCF and how many simulations are rejected due to high founder relatedness, multiple iterations of Ped-sim may be run.

We also sought to identify Gimelfarb (1981)’s Type III relationships, such as double-cousins. We used our simulation pipeline to simulate various Type I and Type II relationship pairs, as well as several Type III relationships. For example, we use “2nd+” degree to refer to second degree relatives who have an additional close relationship through the other parent, e.g., paternal half-siblings and maternal first-cousins. Among the Type III relationships that we simulated are (in order or increasing kinship): double half-cousins (4th+), first-cousin/half-cousins (3rd+), double 3rd degree (Do3rd; e.g., double cousins), and half-siblings/half-cousins and half-siblings/first-cousins (we lump both of these together as 2nd+).

### PONDEROSA’s relationship classifiers

PONDEROSA uses multi-class linear discriminant analysis (LDA) classifiers (Rao, 1948; Fisher, 1936) as implemented in the Scikit-learn Python package (Pedregosa et al., 2011). In short, multi-class LDA attempts to find a linear transformation *ϕ* that projects the data onto a low-dimensional space, maximizing the ratio of between-class scattering and within-class scattering (Li et al., 2006). Labeled data are required to find *ϕ* and unlabeled data can be classified by transforming the data with *ϕ* and taking the classification as the class whose mean vector is closest (in terms of Euclidean distance) to the transformed data point.

LDA assumes that the data are multivariate Gaussian with equal covariance matrices. We find that while feature data in PONDEROSA may not strictly fit those expectations, the LDA classifier still performs better than classifiers that do not make assumptions about the distribution of the data (e.g., linear SVM) (Boser et al., 1992). Lachenbruch and Goldstein (1979) note that LDA may be robust to normality violations (1) when skewness and kurtosis are not extreme and (2) differences of covariance matrices when class means are significantly different, which we observe here. An advantage of LDA over a standard Gaussian mixture model (GMM) is that it avoids the identifiability problem of GMM components. As a note on time complexity, LDA has an *𝒪* (*abc* + *c*^2^) time complexity where *a* is the number of samples, *b* is the number of features, and *c* = *min*(*a, b*) (Cai et al., 2008). For all classifiers in PONDEROSA, *b* = 2, and since there will always be more samples than features, *c ≤* 2. Therefore, the time complexity of the PONDEROSA pipeline can be approximated as linear with the number of pairs. PONDEROSA trains three different LDA classifiers. We first describe the generation of training data, then outline the three LDA classifiers, and end by describing PONDEROSA’s relationship-likelihood tree.

#### Degree of relatedness classifier

The task of inferring the degree of relatedness of a pair is standard and trivial for close relatives in outbred populations; see Ramstetter et al. (2017). However, for relative pairs from populations that exhibit background IBD sharing, the task of inferring the degree of relatedness is less straightforward. For example, Ramstetter et al. (2017) find that KING classifies 1.88% of true 3rd degree relatives from the San Antonio Mexican American Family Study as 2nd degree relatives; in the Himba, we find that KING classifies 21.9% of 3rd degree relatives as 2nd degree relatives. The feature inputs into our degree classifier are the proportions of the genome individuals share IBD1 and IBD2: 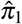 and 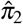, respectively. An graphical example of this classifier is found in Figure 2A (left).

**Figure 2.**
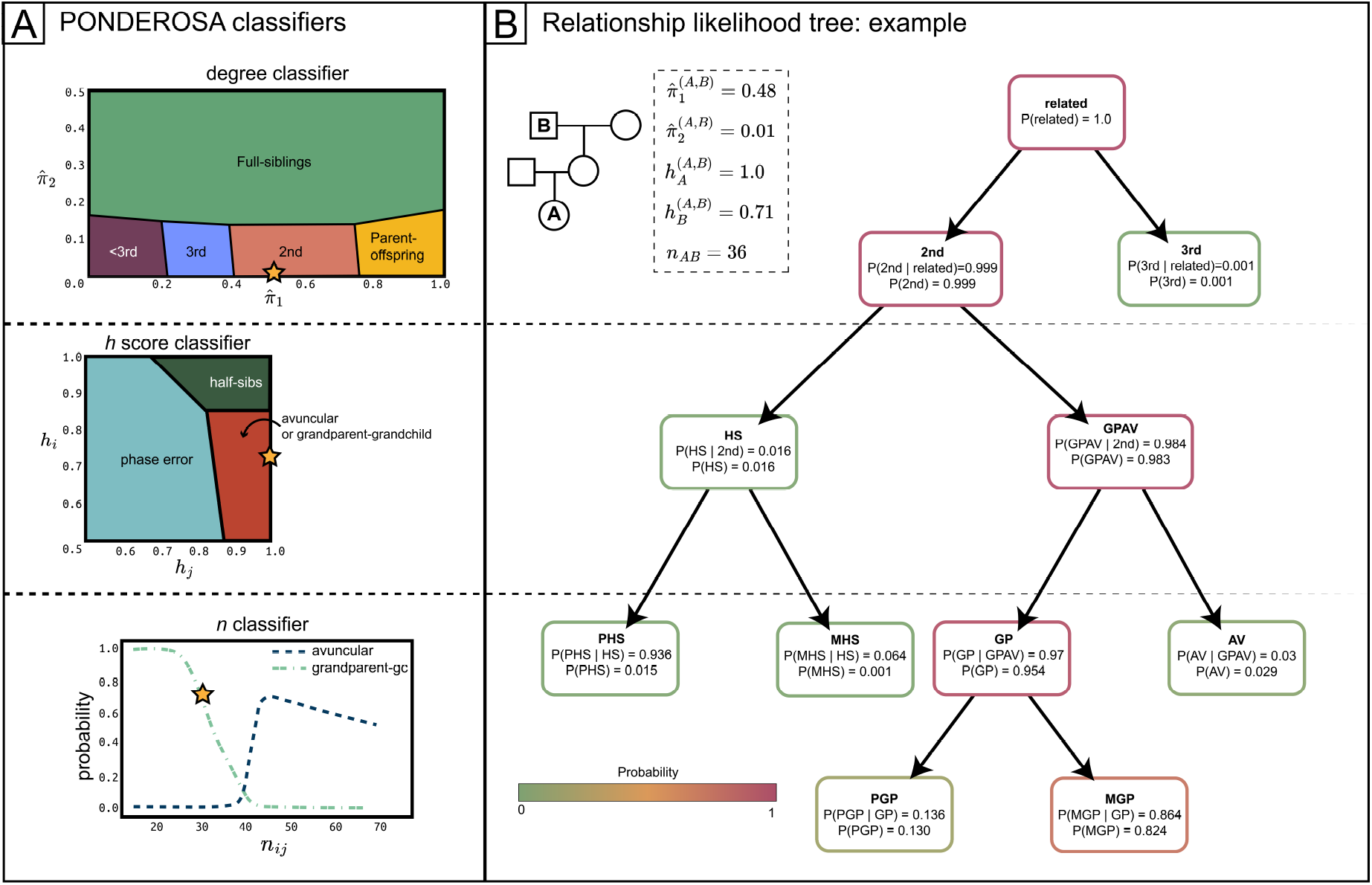
**A** The three classifiers used by PONDEROSA. The star on each plot indicates where an example pair (a maternal grandparent-grandchild) falls on the classifier. The degree classifier predicts the degree of relatedness of the pair based on the proportion of the genome shared IBD1 and IBD2 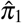 and 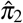, respectively). The plot is colored by the most likely degree of relatedness, but the classifier also outputs class probabilities. The haplotype score classifier predicts whether are second-degree pair is a half-sibling pair or a grandparent-grandchild/avuncular; there is also a third possible classification called “Phase error”, which indicates that a pair’s phase error is too high to reliably distinguish half-siblings from grandparent-grandchild/avuncular. The *n* classifier takes as input the number of IBD segments and total proportion of the genome covered by IBD. Here we are showing the probability of avuncular and grandparent-grandchild as a function of the number of IBD segments shared (probabilities of other relationships not shown). **B** shows an example of the relationship likelihood tree, which is a directed graph, for the same example pair in **A**. Each node is a relationship and child nodes are more specific descriptors of the parent node, e.g., grandparent-grandchild/avuncular (GPAV) is the parent node of grandparent-grandchild (GP) and avuncular (AV). Other nodes include PHS (paternal half-siblings), MHS (maternal half-siblings), PGP (paternal grandparent-grandchild), and MGP (maternal grandparent-grandchild). Each node has a conditional probability, that is the probability of the relationship given that its parent relationship P(child|parent), and a posterior probability P(child). The most likely relationship is a maternal grandparent-grandchild, but if we wanted to be >95% confidence, PONDEROSA would output the most likely relationship as non-sex specific grandparent-grandchild. If we had other information that leads us to believe the pair is actually a half-sibling pair, we could look at the conditional probabilities and find that the pair is most likely a paternal half-sibling pair.

#### Haplotype score classifier

A key difference in IBD sharing between half-sibling pairs and grandparent-grandchild or avuncular pairs is in the distribution of IBD segments across the parental haplotypes. Because half-sibling pairs are related through only a single parent, all the IBD segments that they share should be on the same haplotype, i.e., the paternal haplotype for paternal half-siblings. However, in a grandparent-grandchild/avuncular pair, the genetically younger individual (the grandchild or the niece/nephew) should share all IBD on the same haplotype, but the genetically older individual (grandparent or aunt/uncle) should share the IBD across both parental haplotypes. We developed the haplotype score *h* to describe the IBD sharing on different haplotypes. More specifically, the haplotype score for individual *i* who is related to individual *j* is given by 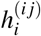. Let *S* be the set of IBD segments shared by *i* and *j*. The length of segment *s* is given by *l*_*s*_ and the haplotype index of individual *i* given by *i*_*s*_. We define the haplotype score as:

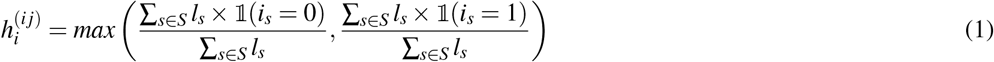

In practice, the haplotype index is arbitrary across chromosomes, so we have to modify the haplotype score as follows:

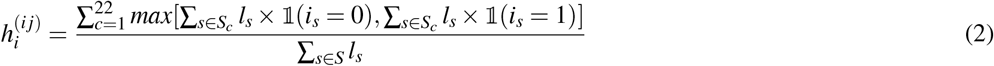

where *S*_*c*_ is the set of IBD segments on chromosome *c*. Equation 2 is equation 1 computed for each chromosome, weighted by the length of IBD on the chromosome.

The haplotype score is sensitive to long-range phasing errors. To address this, we simulated switch errors (SEs) as a Poisson process where the distance between switch errors is *∼* Exp(*d*), where *d* is the mean distance in cM between SEs. Increasing *d* decreases the long-range phase quality; we chose *d* = 25, which approximated a threshold at which the classifier no longer has power to distinguish half-siblings from avuncular/grandparent-grandchild. Next we move along the chromosome. When a SE is encountered, we switch the haplotype index of all IBD segments downstream the SE. If the SE falls in the middle of an IBD segment, the IBD segment is split and the haplotype index of the new downstream IBD segment is switched. If a newly created IBD segment is < 2.5 cM, it is discarded. We then compute the haplotype score using Equation 2. In datasets that do not have the sufficient long-range phase quality, we expect both 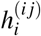 and 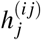to be closer to 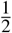.

When phase error is high, we do not wish to make an inference with the haplotype score classifier. Therefore, the classifier has an additional “Phase error” class that is trained on the haplotype scores computed with simulated SEs. If a pair are classified as “Phase error”, the probabilities from the haplotype score classifier will be ignored. Otherwise, the classifier will give the probabilities of the pair being half-siblings or grandparent-grandchild/avuncular. In practice, we re-order the pair such that 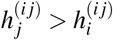, where relative *i* is the grandparent/uncle/niece if the pair are classified as grandparent-grandchild/avuncular. A graphical representation of the classifier can be found in Figure 2A (center).

#### Number of IBD segments classifier

The other statistic we use in PONDEROSA to distinguish among 2nd degree relatives is the number of IBD segments shared the pair (*i, j*), denoted *n*_*i j*_. *n* is not simply number of IBD segments reported from an IBD caller, which would be sensitive to phasing errors and would artificially inflate *n* for grandparent-grandchild and avuncular pairs even in the absence of phase errors. As an example, consider an IBD segment transmitted from grandparent to grandchild; this IBD segment may be a mosaic of the grandparent’s paternal and maternal chromosome. Consequently, an IBD caller would output this as two IBD segments, even though the segment is passed down to the grandchild as a single IBD segment. To compute *n*, we merge IBD segments that are separated by a gap < 1 cM. Importantly, the computation of *n* is agnostic to the haplotype indices of the IBD segments and therefore robust to long-range phasing errors.

Previous work has shown that *n* has power in distinguishing 2nd degree relatives (Henn et al., 2012; Hill and White, 2013; Jewett et al., 2021). Henn et al. (2012) showed that grandparent-grandchildren share fewer IBD segments (but of longer length) than avuncular pairs, as grandparent-grandchildren are separated by two meioses and avuncular pairs are separated by three meioses. We also find that *n* can distinguish maternal half-siblings from paternal half-siblings and maternal grandparent-grandchild from paternal grandparent-grandchild, with maternally-related pairs sharing more IBD segments than paternally-related pairs. This is consistent with previous findings and the higher recombination rate in females (Kong et al., 2002; Caballero et al., 2019; Qiao et al., 2021). In addition to *n*, we include another statistic as training data: 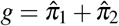, the proportion of the genome covered by IBD to help control for increased *n*, as we might expect that avuncular pairs, for example, that share less IBD overall (due to the randomness of meiosis) might have fewer IBD segments shared than avuncular pairs who share more IBD overall. In Figure 2A (right) we show the probability of avuncular and grandparent-grandchild at a fixed *g* for different values of *n*.

### Relationship likelihood tree

We store the probabilities from the classifiers outlined above in a directed acyclic graph we call the “relationship likelihood tree” (Figure 2B). Each node describes a relationship and child nodes are more specific relationship descriptors than parent nodes, such that the leaf nodes are the most specific pedigree relationship. First, we assign a conditional probability for each node that is the probability of the relationship *given* the pair is the more general parent node relationship, i.e., *P*(child | parent). We next traverse through the tree in a breadth-first manner computing posterior probabilities such that the probability of a child node is given by *P*(child) = *P*(child | parent) *×P*(parent).

There are four primary benefits of the likelihood tree. First, the three classifiers outlined above all output probabilities of various levels, or specificity, of relationships, and the relationship likelihood tree allows for these probabilities to be stored at different levels. Second, the visualization of the tree that we provide as an output is helpful for by-hand pedigree construction, allowing visual access to alternative relationships and their probabilities. Third, the tree easily allows for integration of non-genetic likelihoods, such as likelihoods due to age (as done in Jewett et al. (2021)). Fourth, the data structure allows us to use a simple tree traversal algorithm to output a single most likely relationship depending on our desired minimum probability. For example, setting the minimum probability to 0 will output the most likely leaf node (most specific relationship), but the probability may be small. Alternatively, if we set the minimum probability to 0.80, PONDEROSA will output the most specific relationship with > 0.80 probability.

### PONDEROSA implementation

PONDEROSA takes as input IBD segment estimates and a PLINK-formatted FAM file (to define known paths through the pedigree). Optionally, PONDEROSA accepts KING’s –ibdseg output file, which estimates 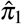 and 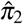 (Manichaikul et al., 2010). We find that KING’s estimates correlate strongly with the estimates computed from the IBD segments themselves (this computation is slow). The user can also supply reported age data (to constrict possible pedigree relationships). PONDEROSA is written in Python 3.10.9 to take advantage of the scikit-learn library for classification (Pedregosa et al., 2011) and can be run on the command line. We designed a data structure to store the pairwise relationship likelihood trees; the data structure contains functions to update probabilities and recompute posterior probabilities. PONDEROSA writes this data structure to a binary file, allowing users to reload the data structure post-hoc, while including non-genetic likelihoods, such as age data, external pedigree information, or population-specific relationship priors (e.g., in the UK Biobank we may expect few, if any, grandparent-grandchildren).

### Data applications

We test the performance of PONDEROSA on 678 Himba individuals genotyped on Illumina H3Africa and MEGAex arrays (Scelza et al., 2020, 2019). The Himba are an endogamous cattle pastoralist population from northern Namibia/southern Angola numbering approximately 50,000 individuals (Scelza, 2011). The Himba diverged from the Herero population in the mid-19th century, likely accompanied by a population bottleneck, and are distinguished today by a variety of cultural norms (Bollig, 1997; Malan, 1995). DNA from Himba families was initially collected to address hypotheses about the frequency of extra-pair paternity. Thus, the study design resulted in a large number of half-sibling relationships. We relied on genetic inference of parent-offspring relationships rather the self-reported assertions. After merging the SNP array datasets and QC (Hardy-Weinberg equilibrium, minor allele frequency, and genotype missingness filters), we subsequently analyzed 486,000 autosomal SNPs. We also test the performance of PONDEROSA using data from BAGS. In BAGS, families were ascertained via asthmatic probands from Barbados through referrals from local clinics/hospitals, and their nuclear and extended family members were recruited into a family-based asthma genetics study (Barnes et al., 1996). As previously described, samples were whole genome sequenced on the Illumina HiSeq 2000, yielding 8,134,080 autosomal SNPs for 981 individuals after QC (MAF 0.05, HWE 1e-6). The data were phased with SHAPEIT2 with the –noped flag and without the –duohmm flag, which would constrain possible haplotypes (O’Connell et al., 2016). KING v2.1.5 was used to find parent-offspring pairs and IBD proportions (IBD1 and IBD2) (Manichaikul et al., 2010). We estimated IBD segments using *phasedibd* (Freyman et al., 2021), setting *use_phase_correction* to False. Otherwise, the resulting IBD segments’ haplotype information would be lost.

We also use a dataset we refer to as the “synthetic Quebec” dataset. This dataset comes from Anderson-Trocmé et al. (2023), who simulated diploid genomes with back-and-forth recombination using *msprime* (Baumdicker et al., 2022). The genomes were simulated through the BALSAC genealogy (Vézina and Bournival, 2020), which contains 6.3+ million individuals dating back to as far as 1680; the genealogy is considered to be a “quasi-complete” genealogy of Quebec (Labuda et al., 2022). To construct our synthetic Quebec dataset, we used the tree sequences provided by (Anderson-Trocmé et al., 2023) and took a random sample of 10,000 individuals from the current generation (individuals in which the metadata indicated that time=0). We simulated mutations at a rate of 9.4 *×* 10^*−*8^ using the Jukes & Cantor ‘69 model that is default with *msprime*. We subset to bi-allelic site with minor allele frequencies > 0.01 and then further randomly downsampled 3% of these sites, which left us with a total of 1,053,687 autosomal sites. We output the resulting tree sequences to VCFs and called IBD using *hap-ibd* (Zhou et al., 2020). We also ran KING, which inferred 231 second degree relative pairs in the 10,000-person subset. As we do not have the true genealogical relationship of the pairs, we did not assess PONDEROSA’s performance on the synthetic Quebec dataset, but used it to demonstrate PONDEROSA’s ability to identify complex, reticulated relationships, Gimelfarb (1981)’s Type III relationships.

## RESULTS

### Overview of PONDEROSA

PONDEROSA is a flexible framework for finding and inferring genetic relatives. Our algorithm infers pairwise degrees of relatedness and, for second degree relatives, infers the specific pedigree relationship of the pair. Unlike other software packages (Jewett et al., 2021; Staples et al., 2014; Qiao et al., 2021), PONDEROSA infers pairwise relationships using IBD sharing data private to each pair. PONDEROSA does not use external relatives or larger pedigree structures to infer relationships. Instead, PONDEROSA uses various summaries of pairwise IBD sharing to train a series of classifiers. The probabilities from these classifiers are stored in what we call a “relationship-likelihood tree”, which is a rooted, directed graph where each node is a relationship label (e.g., third degree relative or paternal half-siblings) and a relationship node’s child nodes are more specific relationships (e.g., the half-sibling node has two child nodes: maternal half-siblings and paternal half-siblings). There are several advantages of storing the probabilities in this format (see Methods).

A major benefit of PONDEROSA is that it trains its classifiers on data computed from known relative pairs in the dataset, which results in performance boosts for datasets from populations with increased IBD sharing due to endogamy or founder events. Other software packages use pre-trained classifiers or otherwise fix model parameters (Jewett et al., 2021; Staples et al., 2014). The most straightforward way that PONDEROSA trains itself is by mining the input data for known relationships, which it finds by tracing parent-offspring lineages. In datasets where PONDEROSA cannot find enough training pairs, we provide a simulation pipeline that uses Ped-sim (Caballero et al., 2019) and the dataset’s genotype data to generate realistic IBD segments for simulated relatives.

### Benchmarking PONDEROSA

In order to evaluate the accuracy of assigning relationships to both first through fourth degree kinship categories and among different types of second degree relationships, we benchmarked the performance of PONDEROSA against publicly available software. We evaluated accuracy among a subset of pairs whose relationship is known either through high-confidence parent-offspring relationship calls or from self-reported pedigree information. Accuracy metrics were tested in two different populations in order to assess the relative effect of endogamy (elevated background IBD) and phase error. We further evaluated the accuracy of the haplotype statistic to correctly orient generational age, again using known ages from the two cohorts as a truth set.

We benchmark the performance of PONDEROSA against ERSA (v2.1) and KING (v2.1.5) (Li et al., 2014; Manichaikul et al., 2010). ERSA was selected because it is an IBD segment-based method; models the number of shared ancestors (and so can distinguish grandparent-grandchild, avuncular, and half-siblings); and infers relatedness relative to the population IBD sharing background (Huff et al., 2011). KING is widely-used, particularly in the GWAS literature and infers the degree of relatedness based on estimates of IBD1 and IBD2 sharing. Importantly, KING’s estimates are based on allele frequencies and *not* IBD segment lengths. However, we do find that KING estimates of IBD1 and IBD2 proportions very closely match estimates computed from the IBD segments in the Himba (Pearson’s correlation; *r*=0.998). The default implementation of PONDEROSA infers relatedness up to the fourth degree and so cannot underestimate relatedness for fourth degree relatives (e.g., by inferring that they are fifth degree relatives). We removed fourth degree pairs that KING and ERSA report as more distantly related than fourth degree so as not to penalize these methods for underestimating relatedness.

PONDEROSA outputs a relationship likelihood for each pair. For our benchmarking analyses, we take the most likely relationship out of the relationships of interest (e.g., to benchmark the degree classifier, we take the most likely relationship out of parent-offspring, full-siblings, second, third, and fourth degree relatives as the “inferred” relationship).

### Assigning degrees of relatedness

First, we analyze PONDEROSA’s performance assigning degrees of relatedness in the Himba compared to KING and ERSA. Overall, PONDEROSA has high sensitivity and specificity across all first through fourth degrees of relatedness. KING frequently over-estimates the degree of relatedness and ERSA’s performance varies across the spectrum of relatedness. For instance, ERSA has high sensitivity and specificity for assigning third degree relatives, but greatly underestimates the relatedness of second degree relatives. Figure 3 summarizes the degree of relatedness assignments for all three algorithms in the Himba. For second degree relatives, PONDEROSA and KING have high sensitivities (97.4% and 99.5%, respectively). However, the specificity of PONDEROSA is high but low for KING, such that KING infers nearly 22% of Himba third degree relatives to be second degree relatives. The relative performances of KING and ERSA as similar to their performances in Ramstetter et al. (2017). In general, PONDEROSA has a slight tendency to underestimate relatedness, underestimating 2.6% and 6.6% of second and degree relatives, respectively. We ran the same analysis for BAGS. We did not expect PONDEROSA to greatly outperform KING because the BAGS individuals come from a relatively outbred population. Indeed, performance is similar, with PONDEROSA performing slightly better than KING at identifying 3rd degree relatives (Figure S1)

**Figure 3.**
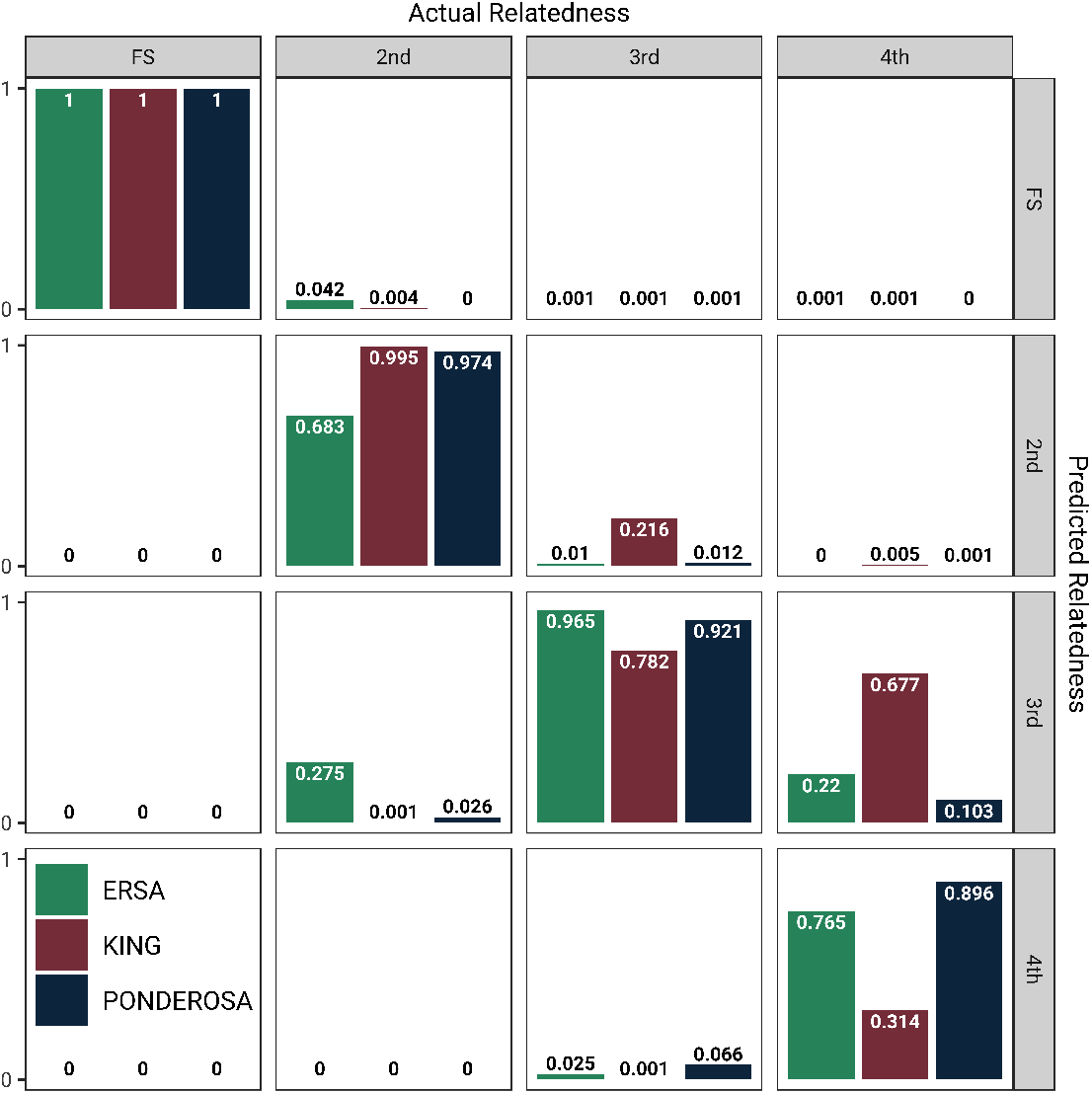
Benchmarking PONDEROSA, ERSA and KING. The bar plots indicate what proportion of actual relatives (columns) are inferred as the row-indicated relationship. For example, KING infers 67.7% of fourth degree relatives as third degree relatives, whereas PONDEROSA infers 10.3% as such. The diagonal indicates the proportion of pairs classified correctly and is equivalent to the sensitivity.

### Classifying types of second degree relationships

Here we compare the performance of PONDEROSA against ERSA in distinguishing half-siblings, grandparent-grandchildren, and avuncular pairs (Figure 4). KING was not included in this analysis because it does not distinguish between specific pedigree relationships. ERSA estimates the number of shared ancestors, which we use to assign a second degree relative: lineal relatives (direct ancestor-descendant relationship such as grandparent-grandchildren) share zero ancestors, half-siblings have one shared, and avuncular have two shared ancestors. In comparing ERSA and PONDEROSA, we ignored PONDEROSA’s sex-inference (for example, aggregating maternal and paternal half-siblings as just half-siblings). PONDEROSA greatly outperforms ERSA across half-sibling, grandparent-grandchild, and avuncular relationships. ERSA commonly misassigns grandparent-grandchild and avuncular relationships as half-siblings, and misassigns more than half of half-siblings as avuncular. We assessed PONDEROSA’s performance in distinguishing maternal/paternal half-siblings and maternal/paternal grandparent-grandchild (Figure S2). For half-siblings, PONDEROSA correctly assigns the sex of the shared parent 87.2% of the time. For grandparent-grandchildren pairs, PONDEROSA assigns 75.9% of pairs as the correct maternal/paternal grandparent.

**Figure 4.**
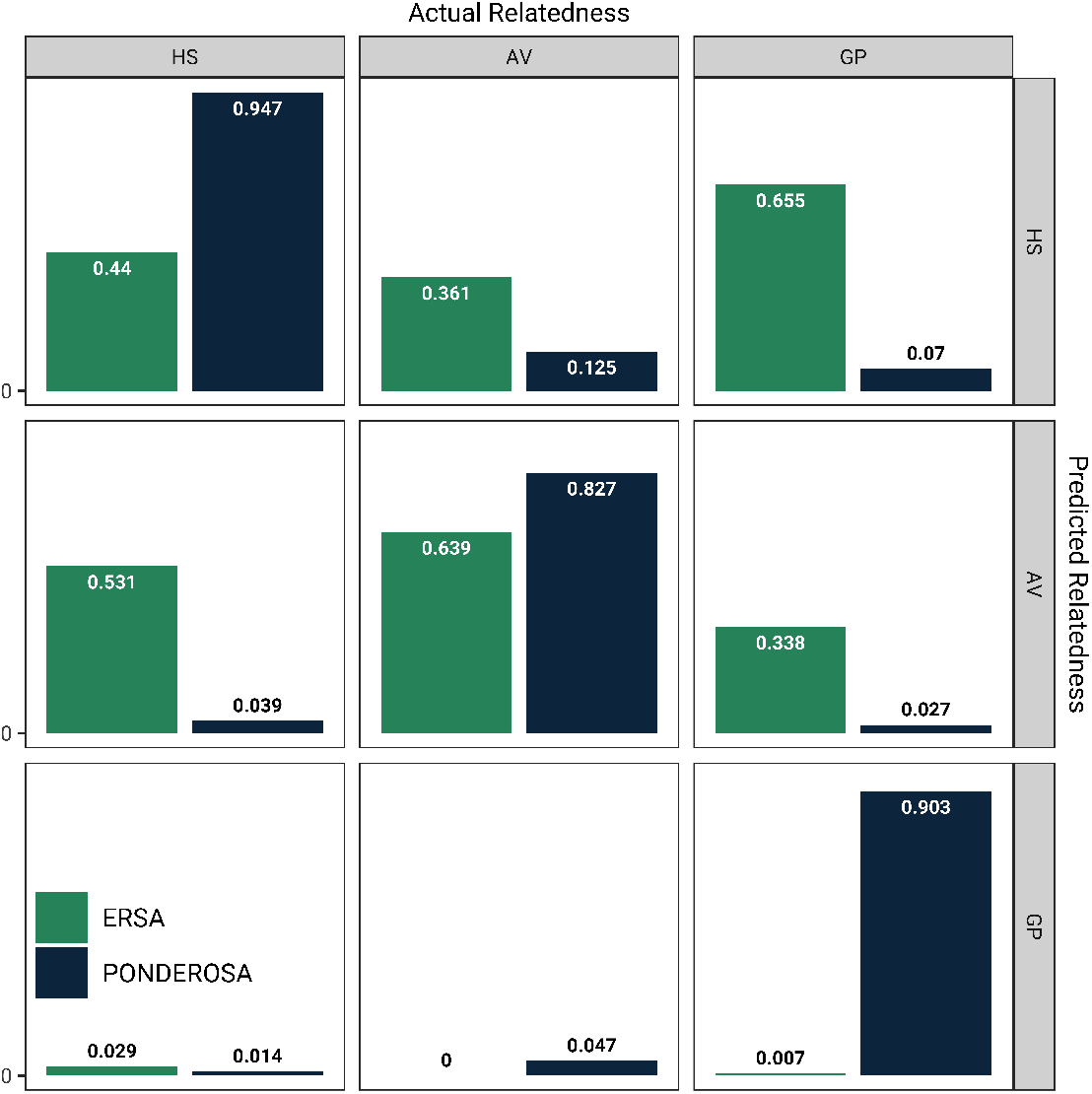
Comparison of ERSA and PONDEROSA assignments of Himba second degree relatives. Sex-specific relationships are aggregated, e.g., half-siblings includes both maternal and paternal half-siblings. PONDEROSA outperforms ERSA in all relationship categories: half-siblings (HS), avuncular (AV), and grandparent-grandchildren (GP).

Next, we looked at the most common pedigree relationship assignments for Himba second degree relatives to assess whether outlying *h* or *n* statistic were driving PONDEROSA’s misclassifications. For half-siblings, the most common misclassification was in the sex of their shared parent (e.g., paternal half-sibling classified as a maternal half-sibling). This suggests that haplotype scores for half-siblings are close to expectation (1) and PONDEROSA performs well in distinguishing half-siblings from avuncular and grandparent-grandchildren. Instead, overlap in distributions of the number of IBD segments *n* for paternal half-siblings and maternal half-siblings drive their misclassification. However, for avuncular and grandparent-grandchildren pairs, the most common misclassification was maternal half-siblings and paternal half-siblings, respectively, suggesting that PONDEROSA’s ability to distinguish avuncular and grandparent-grandchild from half-siblings is limited by high haplotype scores (Equation 2) in the aunts/uncles and grandparents of some avuncular and grandparent-grandchildren pairs.

PONDEROSA also outperforms ERSA in inferring BAGS second degree pairs (Figure 5), but has lower performance than it does in the Himba. This is likely driven by poor phase quality, which makes it more difficult for PONDEROSA to distinguish avuncular from half-siblings, for example, and indeed we see that 38% of BAGS avuncular pairs are misclassified as half-siblings (whereas only 1.5% are misclassified as grandparent-grandchild). We also computed the top-2 accuracy for PONDEROSA’s assignment of BAGS second degree relatives. In Figure S5, we show the top-2 accuracy for the sex-specific second degree relationships (the overall top-2 accuracy is 91.3%). We also computed the top-2 accuracy for non-sex specific second degree relationships, and found an overall top-2 accuracy of 96.6%.

**Figure 5.**
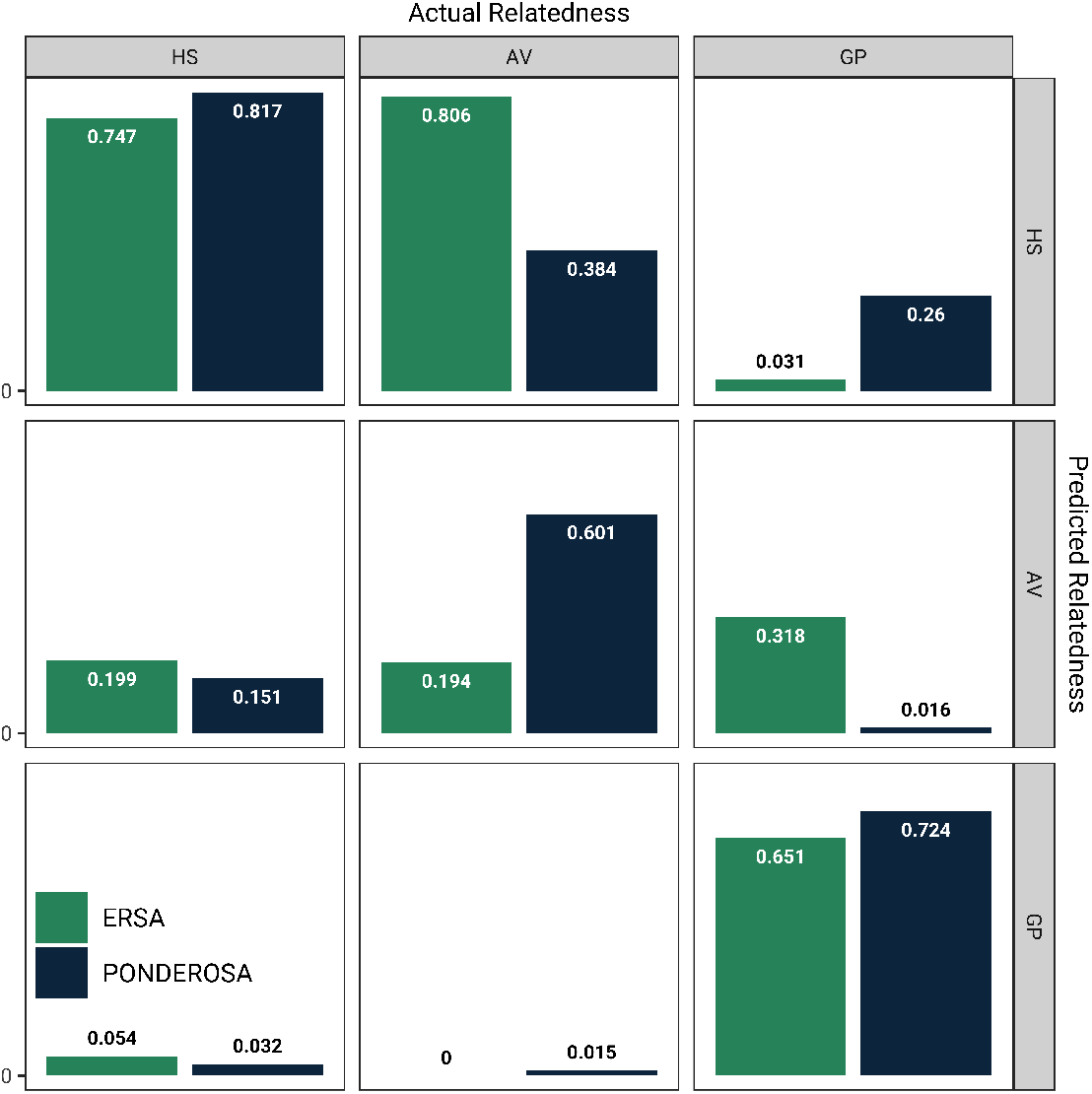
PONDEROSA and ERSA assignments of BAGS second degree relatives. PONDEROSA has lower performance in BAGS, but considerably outperforms ERSA.

We recognize that PONDEROSA’s performance is dependent on high-quality phasing. We tested the robustness of *n* and *h* to long-range phase errors. To do this, we simulated phase errors as a Poisson process parameterized by *λ* = *d*, which is the average distance between switch errors. Increasing *d* increases the frequency of switch errors. We looked at range of *d* = 10 cM to *d* = 200 cM. In Figure S4, we see that *h* is particularly sensitive to long-range phase errors but that *n* is quite robust to phase errors for most values of *d*. To assess the performance of PONDEROSA with poor phase quality, we ran PONDEROSA in the Himba using only the *n* classifier (Figure S3, left). As expected, when only *n* is considered, maternal half-siblings are commonly misclassified as avuncular (and vice versa) and paternal half-siblings are commonly misclassified as maternal grandparent-grandchild. These are pairs of relationships that would otherwise be distinguished with *h*.

### Orienting pairs in a pedigree using haplotype scores

Even when a pair’s pedigree relationship can be accurately assigned, inter-generational pairs pose a challenge in orienting the individuals in a pedigree. Age data can be used but is problematic when it is incomplete, inaccurate or used for orienting non-lineal relationships. For example, an uncle can be younger than his niece. However, if 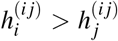, then *i* is the genetically “younger” individual (grandchild/niece/nephew) and *j* is the genetically “older” individual (grandparent/uncle/aunt), as we expect all IBD to fall on the same parental chromosome from the perspective of the genetically younger individual. Using the same logic, individual haplotype scores can be used to orient parent-offspring. For instance, if KING finds an isolated parent-offspring pair and there is no age data, *h* can be used to determine which individual is the parent and which is the offspring.

We assessed the ability of the haplotype scores to correctly orient parent-offspring, avuncular, and grandparent-grandchildren pairs by taking the ratio of the genetically older individual’s *h* and the genetically younger individual’s *h* in known Himba relative pairs (Figure S6). The genetically younger individual should have a higher *h* than the genetically older individual, so this ratio should be less than one. The generational assignment of individuals in pairs that are less than one are correct. In the Himba, haplotype scores correctly orient > 98% of grandparent-grandchild and avuncular pairs, and 99.5% of parent-offspring pairs. Note that most incorrect assignments occur when the difference in haplotype scores is close to zero, suggesting there may be a cutoff where the difference in *h* is too small to accurately orient a pair. The ability to orient parent-offspring pairs depends on the phase quality (see Figure S4) and thus the dataset; in the Himba, the mean difference in *h* is 0.22 but is lower in BAGS (0.17); a lower difference in *h* is more likely to inaccurately orient a parent-offspring pair.

### Identifying Type III relationships

Here we show PONDEROSA’s ability to identify Type III relationships, which are relative pairs who are related through both parents. We simulated these relatives (and other Type I and II relatives) using our simulation pipeline, and ran PONDEROSA using the simulated data as training data. We performed this separately for each of the Himba, BAGS, and synthetic Quebec datasets. We identify several Type III relationships in each population (Figure 6), including 21 3rd+ (e.g., could be first-cousin/half-cousin) in BAGS, four double-cousin equivalents in the synthetic Quebec data, and 197 3rd+ pairs in the Himba. However, we caution that predicting the exact genealogical relationship is difficult. For example, double half-avuncular relative pairs and double-cousins are equivalent in terms of IBD sharing. In the Himba, we observe several double half-avuncular pairs from the known genealogy, but no double-cousins.

**Figure 6.**
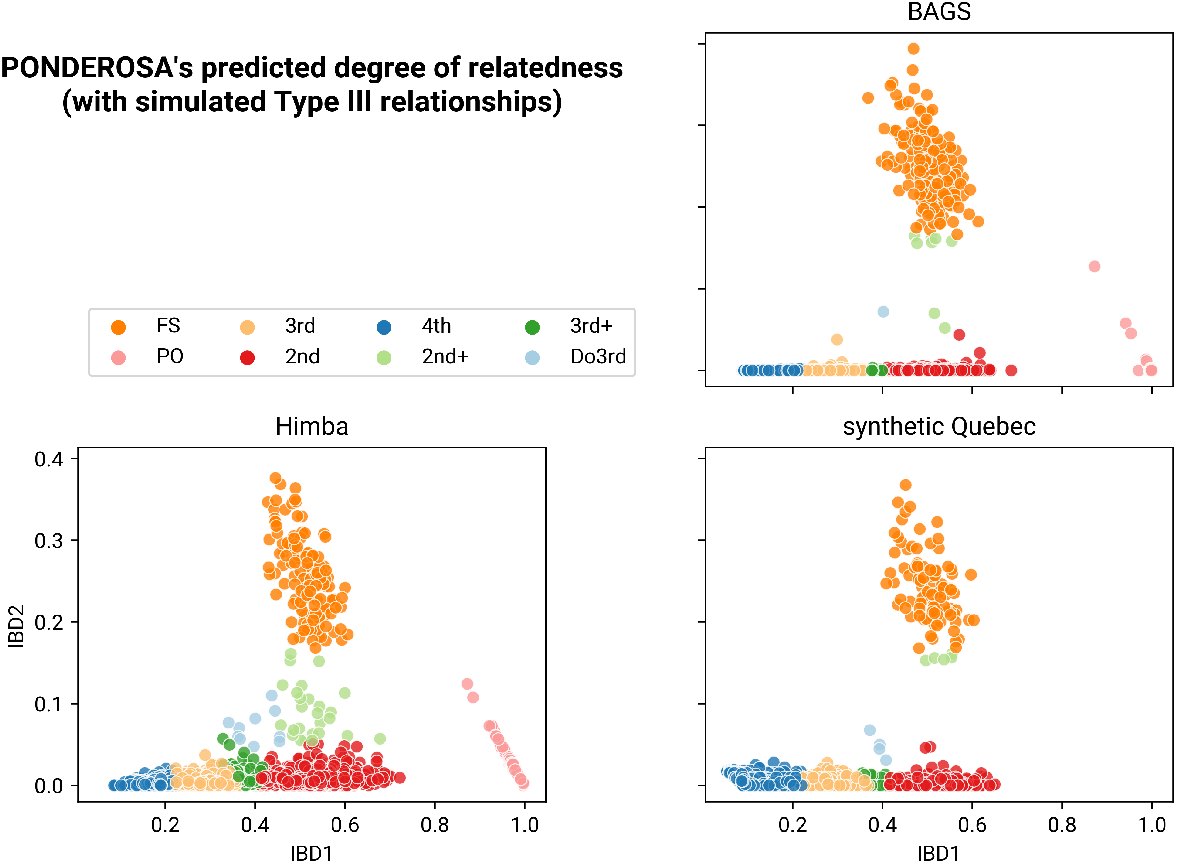
PONDEROSA’s predicted degree of relatedness when modeling type III relationships. For each dataset, we simulated type I, II, and III relationships using the dataset’s genotype data as input. The type III relationships we simulated were double half-cousins (4th+), double 3rd degree (Do3rd), first-cousin/half-cousin (3rd+), and half-sibling/first-cousin (2nd+). These simulated relatives and their IBD segments were used as input training data for PONDEROSA. Here we show PONDEROSA’s predicted degree of relatedness for each dataset’s relative pairs, plotted by their IBD1 and IBD2 proportions. Each population has several type III relationships predicted by PONDEROSA, particularly in the Himba and synthetic Quebec datasets.

## DISCUSSION

PONDEROSA offers a new approach for inferring pedigree relationships from genotype data by initially identifying high-confidence relationships already present in the dataset and using them as a training set for a machine learning classifier. The machine learning steps of PONDEROSA are key to its application in diverse human populations. PONDEROSA trains a linear discriminant analysis (LDA) classifier with the IBD1 and IBD2 values of high-confidence relatives from a given dataset to identify putative second degree relatives. This reduces the danger of assigning a second degree relationship to actual third degree relatives. We then use two additional LDA classifiers to infer the specific pedigree relationship of these putative second degree relatives. One of these classifiers uses the number of IBD segments, which provides resolution, particularly in distinguishing avuncular versus grandparent-grandchild, maternal half-siblings versus paternal half-siblings, and maternal grandparent-grandchild and paternal grandparent-grandchild. The number of IBD segments is not enough for total resolution: in particular, the distributions of the number of IBD segments overlap considerably for both maternal half-siblings and avuncular pairs. To that end, we developed the haplotype score (Equation 2) which we use to distinguish half-siblings from grandparent-grandchild/avuncular when long-range phase quality is high. The haplotype score can also be used to differentiate the genetically younger/older individuals in avuncular, grandparent-grandchild, and parent-offspring pairs when age data is not available. A key innovation of PONDEROSA is the training step, which uses true relative pairs from the dataset such that deviations from an idealized population or technology-specific biases are incorporated into the segment lengths distributions for a relationship pair.

We test our approach on two datasets with different population histories: the endogamous Himba of northern Namibia and an African-descent population from Barbados (BAGS). These datasets are realistic scenarios for PONDEROSA’s best use case: both have dense family structure present with many parent-offspring and full-sibling pairs but also many unresolved close relative pairs. Our results with the Himba show the importance of our machine learning step in inferring degrees of relatedness: 20% of Himba third degree relatives are inferred as second degree relatives by KING. Considering these pairs as second degree would drastically change the pedigree structure and downstream inferences. Additionally, PONDEROSA has high accuracy in inferring paternal half-sibling, maternal half-sibling, and avuncular categories. The accuracy can remain high with few training pairs per category, e.g. accuracy increases only marginally as the number of training pairs increases above 10 per relationship category. For comparison, there are between 300 and 400 training pairs per relationship in the Himba dataset we use. These findings show that PONDEROSA will retain its accuracy even in datasets with few training relationships. PONDEROSA has poorer performance in the BAGS dataset, but has high top-2 accuracy, which is still useful, particularly for by-hand pedigree construction. We also show that PONDEROSA can identify more complex pedigree relationships, such as double-cousins, and demonstrate this feature on the Himba and BAGS dataset, as well as a synthetic Quebec dataset that we generated from publicly-available simulated Quebecois genomes from Anderson-Trocmé et al. (2023).

When the long-range phase quality is high, the haplotype score reflects the how much IBD is shared on a single parental chromosome. However, the haplotype score is sensitive to phasing errors. An indicator of poor phase quality is when both individuals in the pair have low haplotype scores. PONDEROSA’s haplotype score classifier will classify these individuals as having high phase error, and will rely only on the IBD segment number classifier. Another limitation of the haplotype score is that the inter-chromosomal haplotype indexes are arbitrary, such that it is unknown which haplotype corresponds to the same parent between chromosomes. This is the case for all major haplotype phasing algorithms (Browning et al., 2021; Hofmeister et al., 2023). While Equation 1 is the strict definition of the haplotype score, we apply Equation 2 because the inter-chromosomal phase is used. Equation 1 would give us higher power to distinguish grandparent-grandchildren/avuncular from half-siblings. The haplotype score will become more powerful with advances in long-range phasing (Liao et al., 2023; Maestri et al., 2020; Xu and Dixon, 2020), inter-chromosomal phasing (Hofmeister et al., 2021; Noto and Ruiz, 2022; Williams et al., 2024), and as sample sizes get bigger (enabling higher quality phasing). For example, Williams et al. (2024) found that more than half of non-Asian 23andMe trio children have zero long-range switch errors on chromosome 2.

We also created a data structure for storing the probabilities from the three LDA classifiers, called the relationship likelihood tree. We were motivated by the fact that relationships can be called by varying levels of specificity, e.g., half-siblings versus paternal half-siblings, and that probabilities decrease as the specificity of the relationship label increases. We designed PONDEROSA to assist with by-hand pedigree construction, and the likelihood tree allows researchers to visualize the various probabilities. For example, if the most likely relationship inferred by PONDEROSA is a paternal half-sibling, but there is reason to believe that paternal half-siblings are rare, we can easily use the likelihood tree to find the next most likely non-paternal half-sibling relationship.

Overall, our results show the power of considering the haplotype state of IBD segments in inferring pedigree relationships and sex-specific recombination; haplotype state is not currently considered in any publicly available kinship inference algorithms. Advances in phasing will only improve the performance of PONDEROSA. As genetic databases increase in size, there is an increasing need to construct extended pedigrees with members of the databases without reliance on self-reported relationships, which may be either absent, incomplete, or inaccurate. PONDEROSA offers a solution for building extended pedigrees and finding second degree relatives. Building pedigrees out of large genetic datasets will advance the study of disease and recombination across a range of diverse populations.

## AUTHOR CONTRIBUTIONS

C.M.W., B.M.H., and S.R. conceived the algorithm design; B.A.S. collected the DNA samples and pedigree interviews; C.M.W. performed analysis of the data; C.M.W. wrote the algorithm code; R.A.M., H.W., K.C.D. provided genetic data; E.L. provided computational support; N.F.P. and S.D.S. performed tests of the algorithm; C.M.W., C.R.G, S.R. and B.M.H. wrote the main manuscript; All authors have read and agreed to the published version of the manuscript.

## INSTITUTIONAL REVIEW

UC Davis IRB determined that analysis in this manuscript did not constitute human subject research (1767383-1; May 24, 2021). Please see Data availability for institutional reviews associated with the original data collection.

## DATA AVAILABILITY

The genetic datasets used herein are available via dbGaP: phs001995.v1.p1 (Himba) and dbGaP: phs001143.v3.p1 (BAGS). A beta version of PONDEROSA is available: https://github.com/williamscole/PonderosaBeta

## FUNDING

This work was supported NIH grants R35GM133531 (to B.M.H.), R35GM139628 (to S.R.), and T32 GM128596 (C.M.W. and S.R). This work was also funded by the NSF (BCS-1534682 to B.A.S. and GRFP to C.M.W). For the purpose of open access, the author has applied a CC BY public copyright licence to any Author Accepted Manuscript version arising from this submission.

## ACKNOWLEDGMENTS

We thank the community of Omuhonga for their continued support and contribution of genetic data used in this project. J. Jakurama, C. Louis, and K. Ngombe acted as research assistants and translators in Namibia during the original data collection. We thank members of the Henn Lab for beta-testing versions of PONDEROSA and their feedback. Nuria Melisa Morales Garcia designed the logo for PONDEROSA. This research was supported in part by the Division of Intramural Research of NIAID, NIH.

## CONFLICTS OF INTEREST

The authors declare no conflict of interest.

## SUPPLEMENTARY MATERIALS

**Figure S1.**
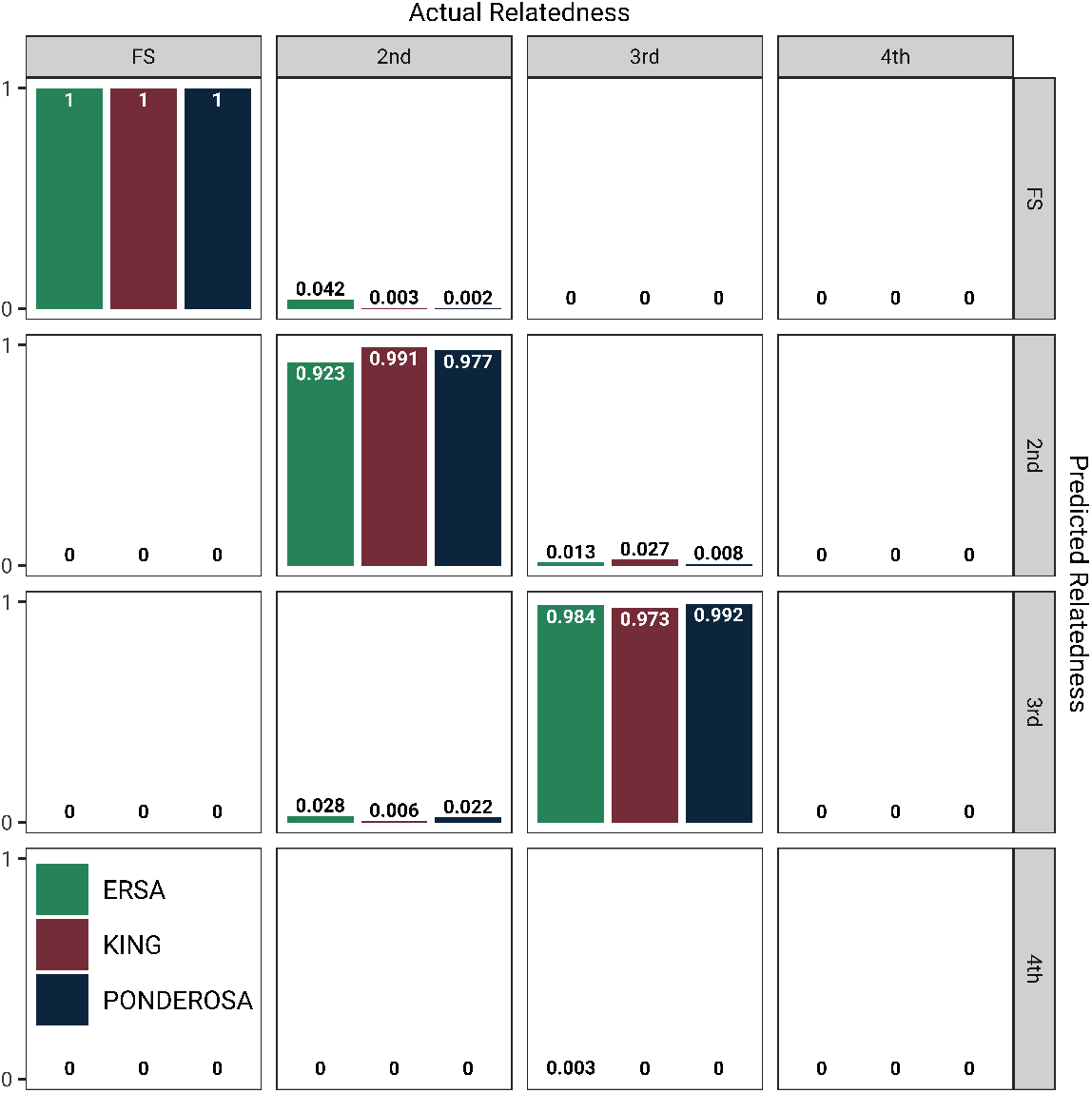
PONDEROSA and KING assignment of degrees of relatedness in BAGS. Because BAGS is relatively outbred, the performance of the two algorithms is similar, although there are slight performance gains in PONDEROSA in assigning 3rd degree relatives.

**Figure S2.**
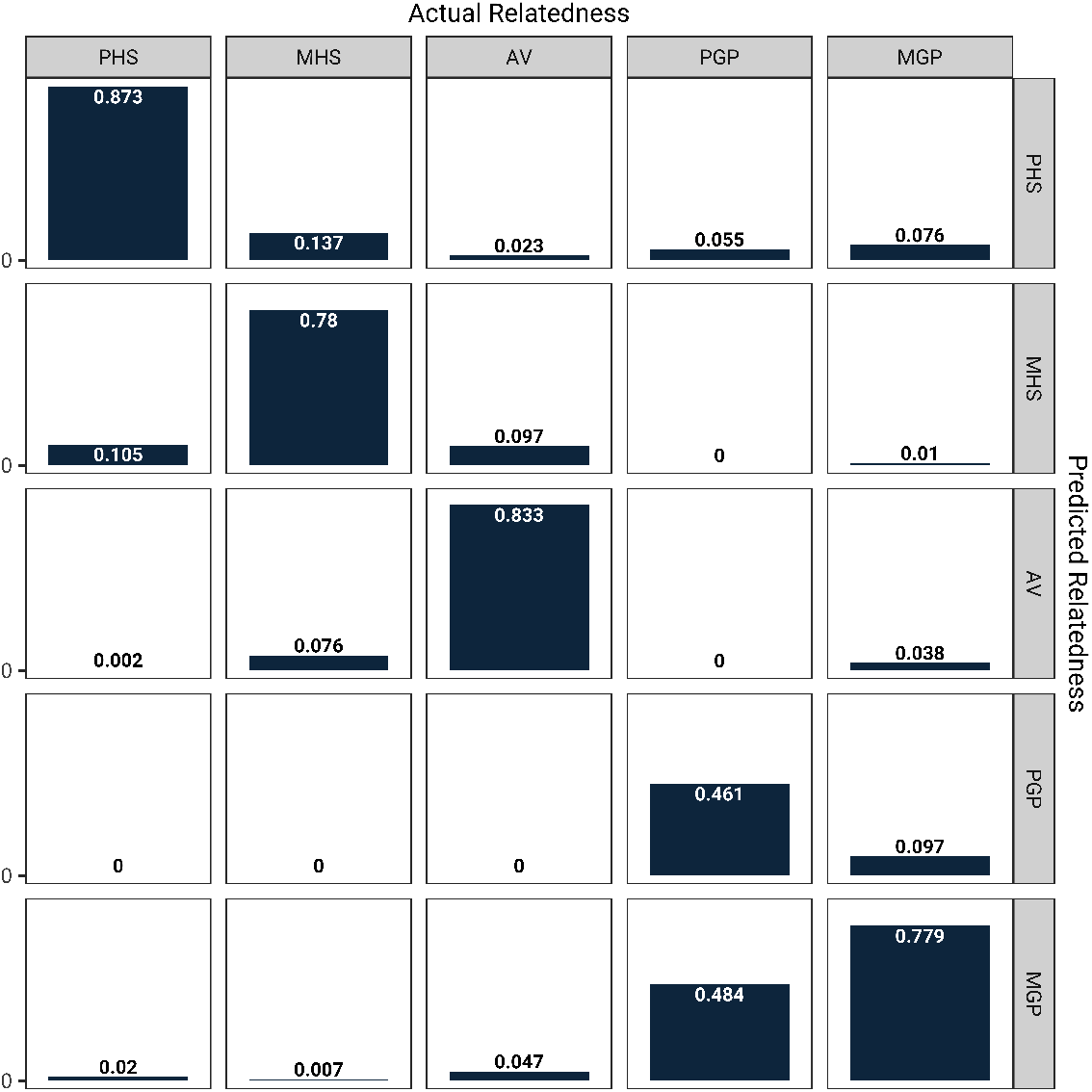
PONDEROSA assignments of Himba second degree relatives with sex-specific relationships. For most relationships, the most commonly misclassification is the sex of the relationship, e.g., 10.5% of paternal half-siblings are inferred as maternal half-siblings.

**Figure S3.**
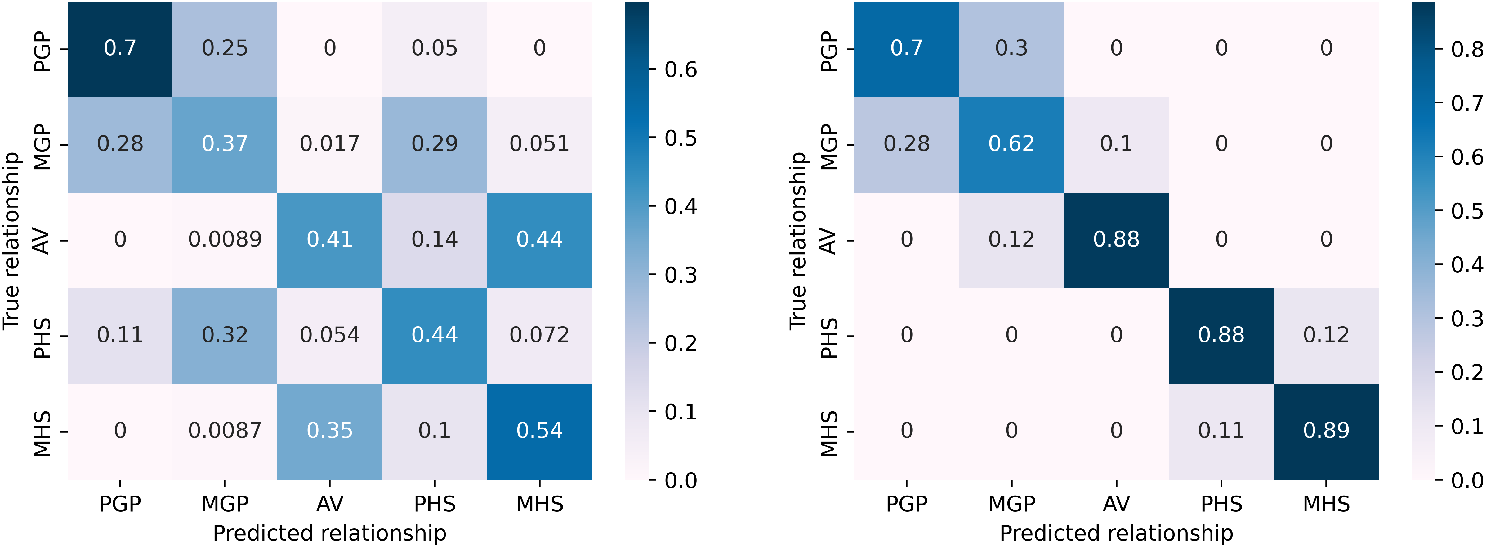
The performance of PONDEROSA without the *h* classifier (left) and with the *h* classifier (right). PONDEROSA performs better with the *h* classifier, but is still informative.

**Figure S4.**
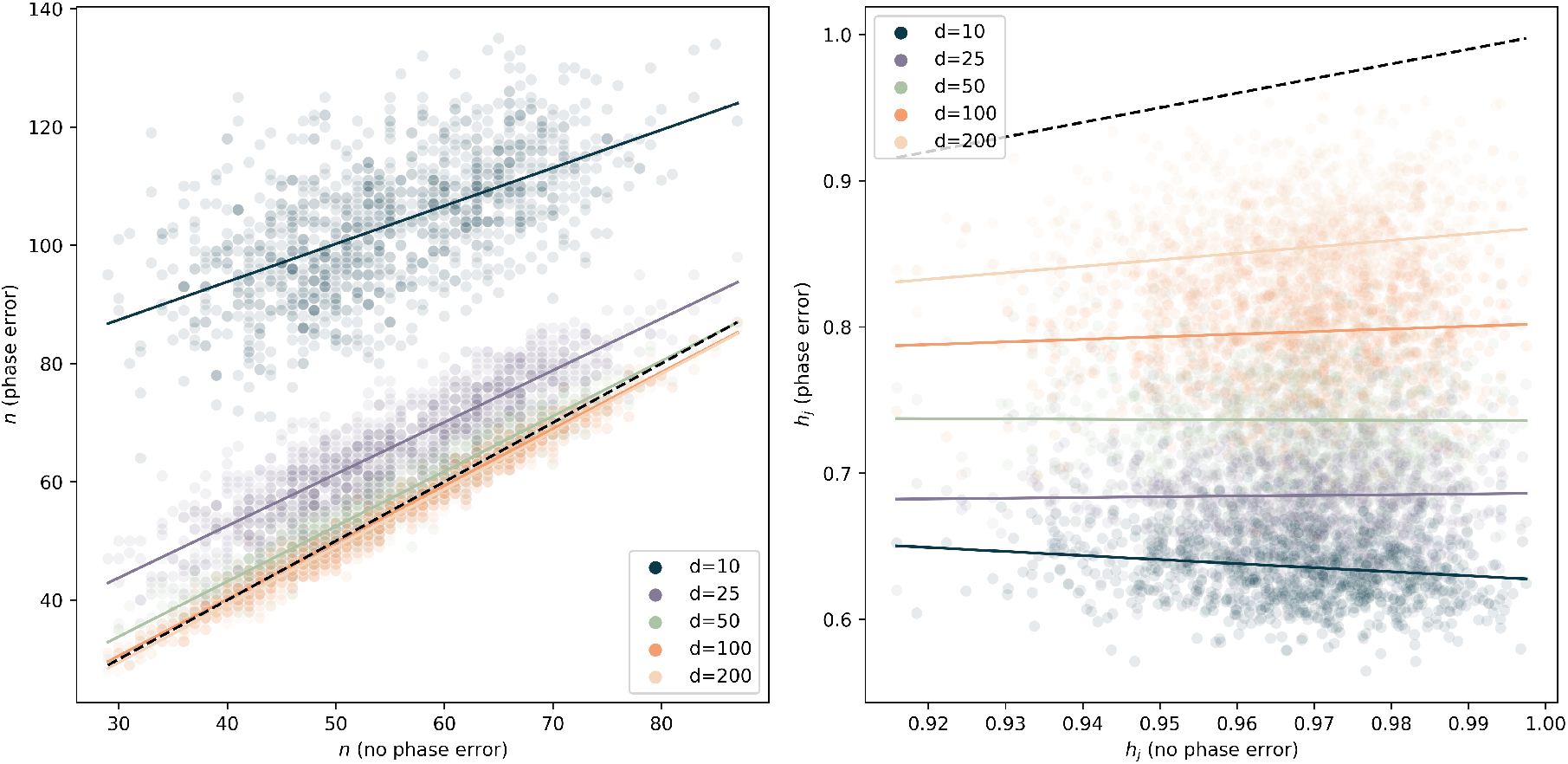
The robustness of the number of IBD segments estimate *n* (left) and the haplotype score *h*_*j*_ of simulated Himba half-siblings (IE(*h*_*j*_) = 1), right. The x-axis is the true value of the statistic and the y-axis is the value of the statistic with phase error (the dashed line is the 1:1 line). We introduce phase errors as a Poisson process with mean *d*, which is the mean distance between haplotype-switching phase errors (lower *d* is lower quality phase). We plot the individual points for each value of *d* and draw a line of best fit through the points.

**Figure S5.**
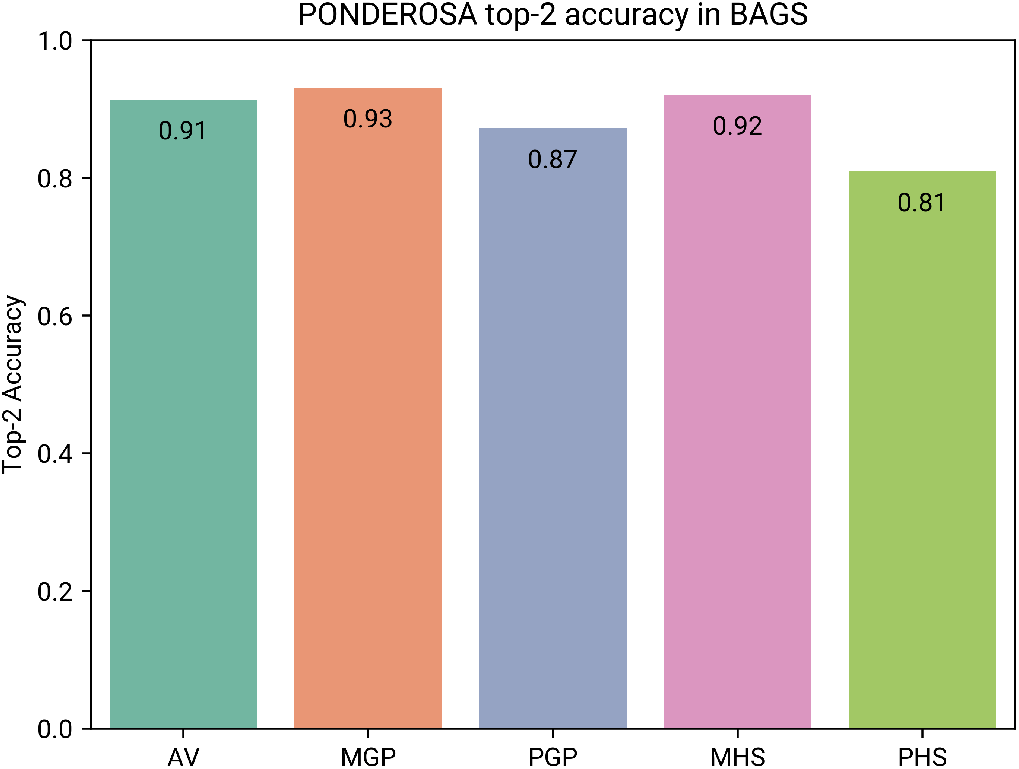
PONDEROSA’s top-2 accuracy in the BAGS dataset. The top-2 accuracy is the proportion of each relationship category’s pairs in which one of the two most probable relationships is the true relationship. Here overall top-2 accuracy is 91.3%, and when we do not consider sex-specific relationships, the overall top-2 accuracy is 96.6%.

**Figure S6.**
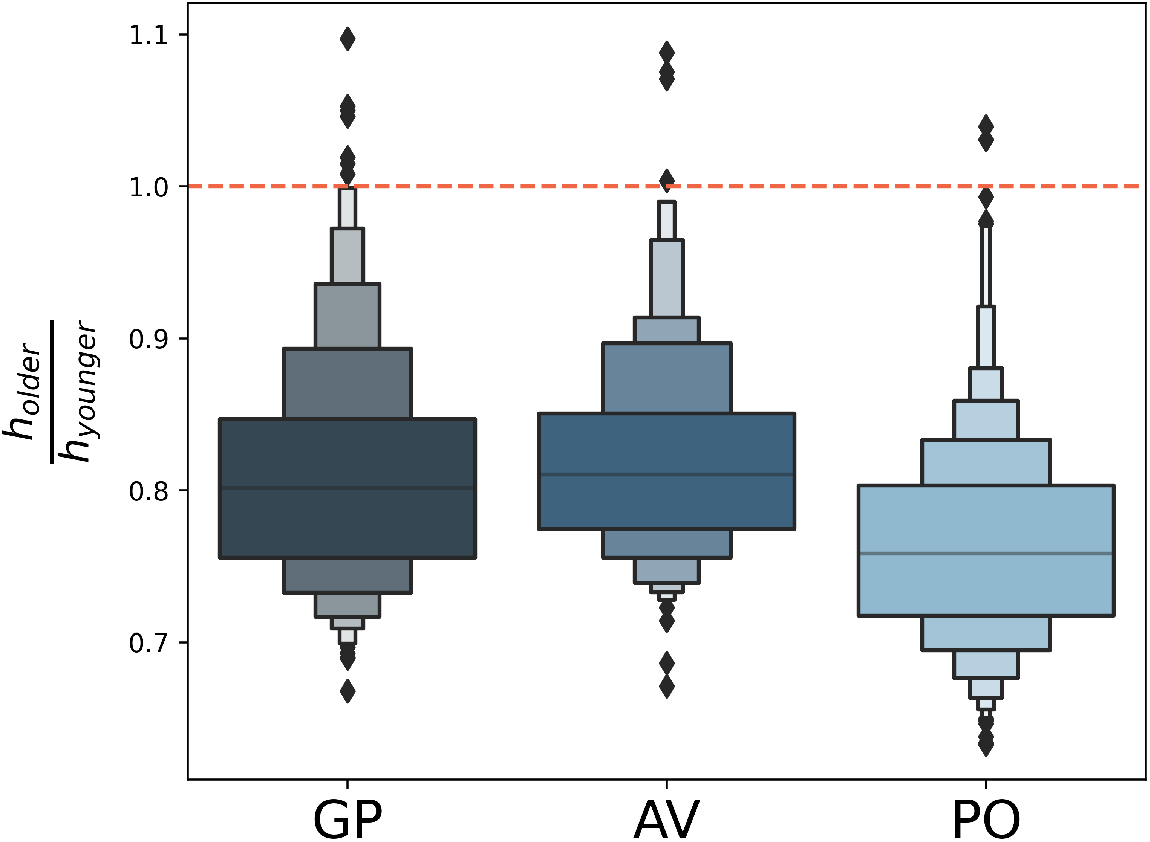
Haplotype score ratios of the genetically older individual’s *h* and the genetically younger individual’s *h*. The expectation is that *h*_*older*_ < *h*_*younger*_, such that the ratios fall below 1. This is true for > 98% of grandparent-grandchild (GP), avuncular (AV), and parent-offspring (PO) pairs, suggesting that the haplotype score is useful for orienting these relationships without the need for age data.

## Notes

https://github.com/williamscole/PONDEROSA

### Competing Interest Statement

The authors have declared no competing interest.

### Summary of Updates

Text has been revised to reflect new features of the algorithm, revised figures and updated author list.

