## Supplemental Figures for "A rapid, accurate approach to inferring pedigrees in endogamous populations"

### SUPPLEMENTARY MATERIALS

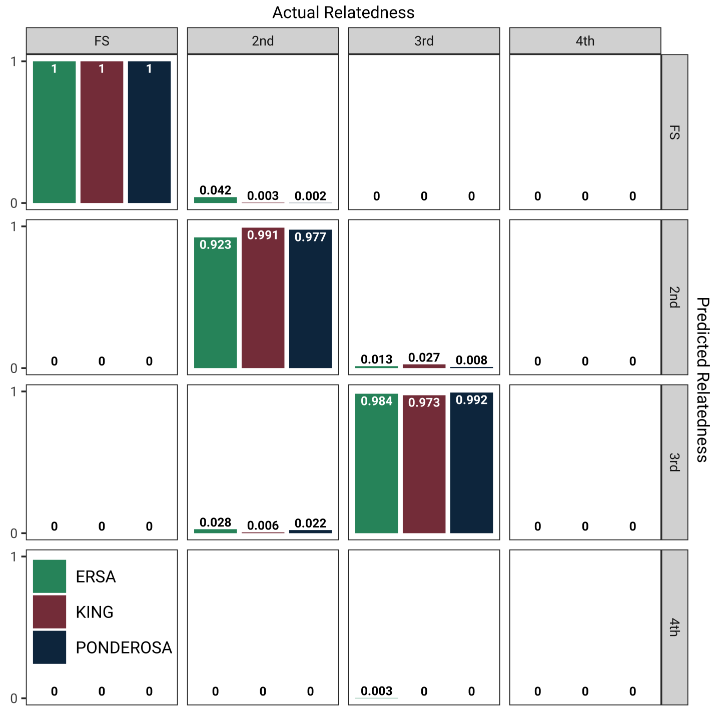


**Figure S1.** PONDEROSA and KING assignment of degrees of relatedness in BAGS. Because BAGS is relatively outbred, the performance of the two algorithms is similar, although there are slight performance gains in PONDEROSA in assigning 3rd degree relatives.

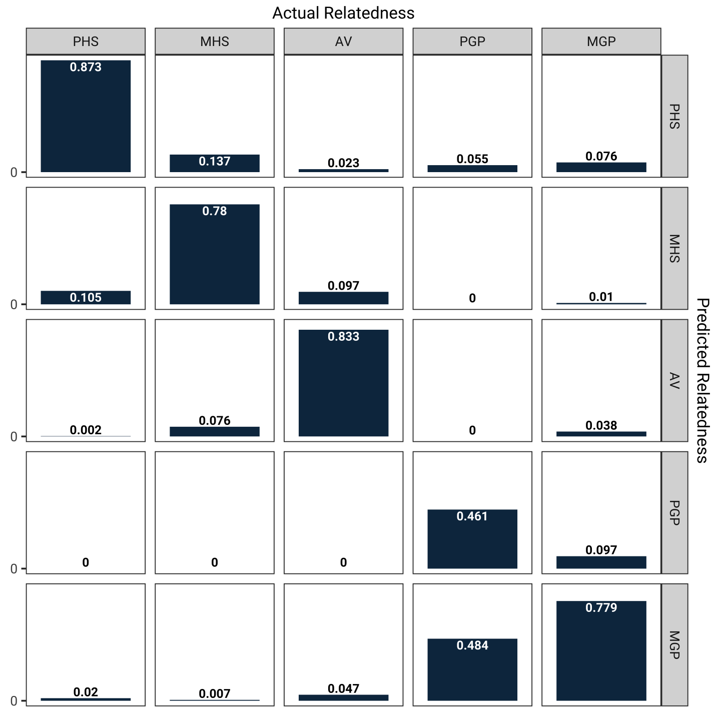


**Figure S2.** PONDEROSA assignments of Himba second degree relatives with sex-specific relationships. For most relationships, the most commonly misclassification is the sex of the relationship, e.g., 10.5% of paternal half-siblings are inferred as maternal half-siblings.

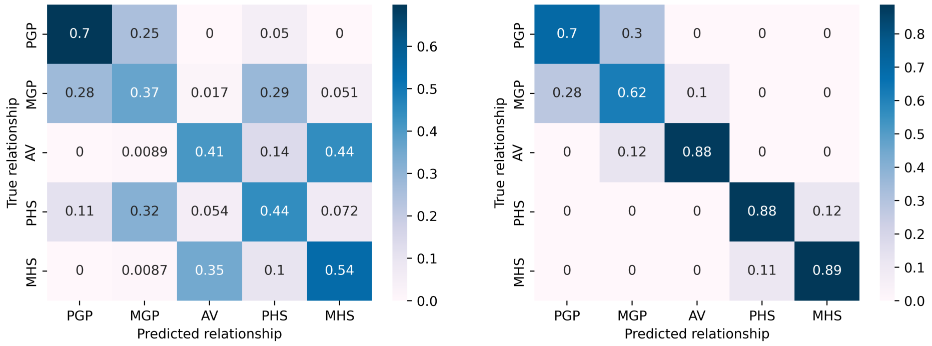


**Figure S3.** The performance of PONDEROSA without the $h$ classifier (left) and with the $h$ classifier (right). PONDEROSA performs better with the $h$ classifier, but is still informative.

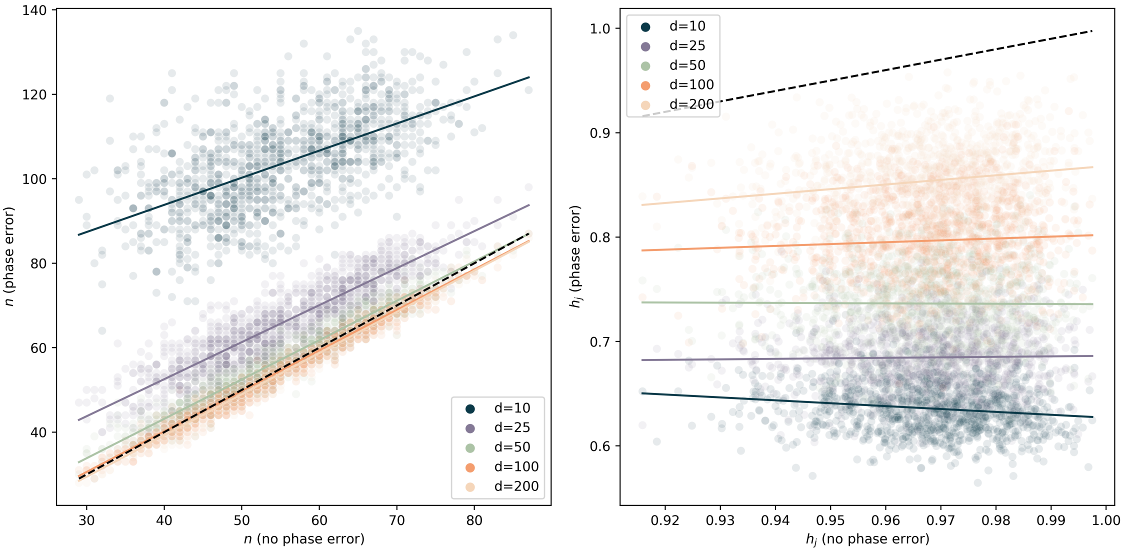


**Figure S4.** The robustness of the number of IBD segments estimate $n$ (left) and the haplotype score $h_j$ of simulated Himba half-siblings ($\mathbb{E}(h_j) = 1$), right. The x-axis is the true value of the statistic and the y-axis is the value of the statistic with phase error (the dashed line is the 1:1 line). We introduce phase errors as a Poisson process with mean $d$, which is the mean distance between haplotype-switching phase errors (lower $d$ is lower quality phase). We plot the individual points for each value of $d$ and draw a line of best fit through the points.

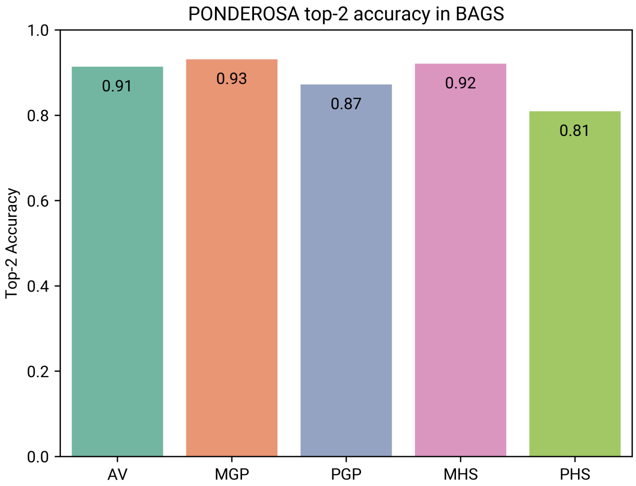


**Figure S5.** PONDEROSA's top-2 accuracy in the BAGS dataset. The top-2 accuracy is the proportion of each relationship category's pairs in which one of the two most probable relationships is the true relationship. Here overall top-2 accuracy is 91.3%, and when we do not consider sex-specific relationships, the overall top-2 accuracy is 96.6%.

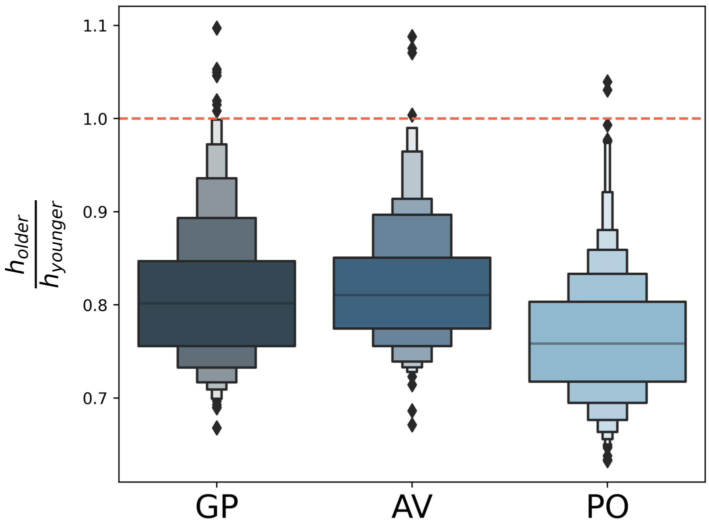


**Figure S6.** Haplotype score ratios of the genetically older individual's $h$ and the genetically younger individual's $h$. The expectation is that $h_{older} < h_{younger}$, such that the ratios fall below 1. This is true for $> 98\%$ of grandparent-grandchild (GP), avuncular (AV), and parent-offspring (PO) pairs, suggesting that the haplotype score is useful for orienting these relationships without the need for age data.
